# An open-source cryo-storage solution

**DOI:** 10.1101/184168

**Authors:** Eveline Ultee, Fred Schenkel, Wen Yang, Susanne Brenzinger, Jamie S. Depelteau, Ariane Briegel

## Abstract

The field of cryo-electron microscopy is a rapidly growing method in structural biology. With this development, access to cryo-EM facilities becomes a bottleneck that results in long wait times between sample preparation and data acquisition. To improve sample storage, we developed a cryo-storage system with a more efficient and larger storage capacity that enables cryo-sample storage in a highly organized manner. This system is simple to use, cost-effective and easily adaptable for any type of grid box and storage dewar and any size cryo-EM laboratory.

---

The field of cryo-electron microscopy (cryo-EM) has expanded over the last few decades and is rapidly growing (Kühlbrandt, 2014; Nogales, 2016). Recent advances in instrumentation (Grigorieff, 2013) and data processing (S. H. W. Scheres, 2012; S. H. w Scheres, 2014) have elevated electron cryotomography and cryo-EM single-particle analysis to the forefront of the study biological structures, both *in vivo* and *in vitro.* Applications range from solving the structures of proteins at angstrom level resolution (Banerjee et al., 2016) to visualizing molecular machines within intact cells (Oikonomou et al., 2016).

While the demand for cryo-EM is expanding rapidly, access to the facilities that provide the necessary instrumentation is limited. The high cost of state-of-the-art cryo microscopes, expert staff and support infrastructure prevents most research groups from obtaining their own core equipment. As such, shared facilities or national centers are becoming the predominant mode of access to cryo-EM for many research groups (Stuart, Subramaniam, & Abrescia, 2016). Unfortunately, the relatively limited access and the ever-rising demand often lead to a long waiting time between cryo-EM sample preparation and data acquisition. Consequently, cryo-EM samples accumulation in either individual research groups or microscopy facilities has been a commonly encountered issue for the entire EM field.

Proper storage of cryo-samples is essential for all cryo-EM laboratories. For cryogenic electron microscopy, biological samples are rapidly frozen by plunging them into a cryogen such as liquid ethane (McDowall et al., 1983) or a mixture of liquefied gasses such as ethane and propane (Tivol, Briegel, & Jensen, 2008). This preserves the native state of the samples within crystal-free vitreous ice. The vitrified samples must be maintained at, or below, 120 K during all subsequent storage, transfer and imaging steps to prevent thawing or the formation of crystalline ice, which will destroy the samples (Tocheva, Li, & Jensen, 2010).

Many labs still store their cryo-EM grid boxes in submerged labeled plastic screw-cap conical tubes attached to strings that hang from the neck of the cryogenic storage dewar. This inexpensive and straightforward ‘tube-with-ropes’ storage system has met the need of many labs, typically in situations where there are few users or where throughput is high and samples need not to be stored for extended periods (Iancu et al., 2007). However, as cryo-EM increases in popularity and the field changes towards multi-user laboratories and facilities, this improvised grid storage method is often no longer suitable. As more plastic tubes are added to a storage dewar, the strings can become entangled, labels can fall off and important samples can be mixed up, lost in the dewar or ruined due to excessive handling, unsecured lids or damaged tubes.

Here, we describe a cryo-storage system with a more efficient and larger storage capacity in the dewar that enables cryo-sample storage in a highly organized manner. This system is simple to use, cost-effective and easily adaptable for any type of grid box and storage dewar and any size cryo-EM laboratory.

## The cryo-storage design

Our cryo-storage utilizes “pucks”; circular metal containers that stack vertically within a dewar canister to optimize available space (figure 1). This design was inspired by a storage system developed by Dr. Alasdair McDowall (University of Queensland, Brisbane, Australia) and later adapted for Grant Jensen’s Laboratory at Caltech (Pasadena, CA, USA), where grid boxes are stored loosely in circular aluminum storage containers. We developed our system for use in the XSS-48/10 model liquid nitrogen storage dewar (VWR systems), which features a 119 mm neck and 10 stainless steel internal canisters (270 mm high by 73 mm in diameter). Each canister has a hook, which suspends it from the lip of the dewar and facilitates its access. The dimensions of the “pucks” and associated parts are easily adaptable for other cryo-storage dewar models.

**Figure 1.**
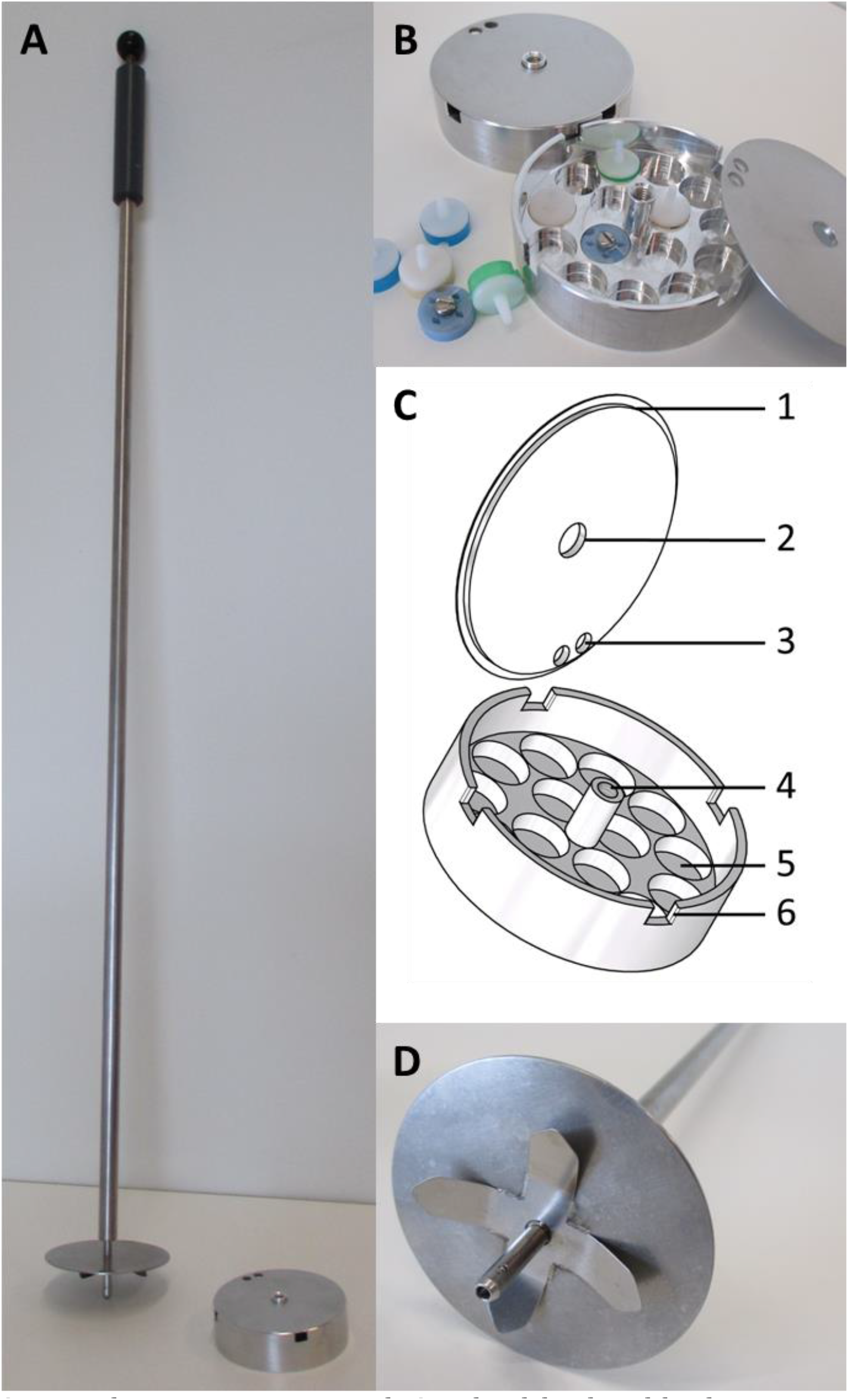
The cryo-storage system designed and developed for the preservation of cryo-EM samples. The cryo-storage system consists of pucks that are vertically stacked in dewar canisters and can be transferred with a retrieval-rod (A). The storage pucks are designed to hold 15 cryo-EM grid boxes (B). The pucks have separate lids with an engraved edge (C-1), two holes to manipulate the lid with tweezers (C-3) and a central hole to (C-2) to fit the central docking cylinder of the puck (C-4). Each puck has grid-boxes sized holders (C-5) and four notches at the edge to enable liquid nitrogen to flow into the puck (C-6). The retrieval-rod is optimized with leaf-springs and a self-locking pin (D).

The circular storage “pucks” for our system are machined from a billet of aluminum 51 ST which is lightweight, a good thermal conductor and easy to machine. The pucks developed for our dewar are 23 mm in height and have a diameter of 70 mm, slightly smaller than the diameter of the canisters they are housed in (figure 1A, 2). The inner bottom surface of each puck is further machined to hold 15 standard grid boxes upright in a single layer, keeping them organized and easy to retrieve (figure 1B, figure 1C–5). Four notches at the rim of the cryo-puck (figure 1C–6) allow the cryo-puck to rapidly refill with liquid nitrogen after insertion into the dewar.

**Figure 2.**
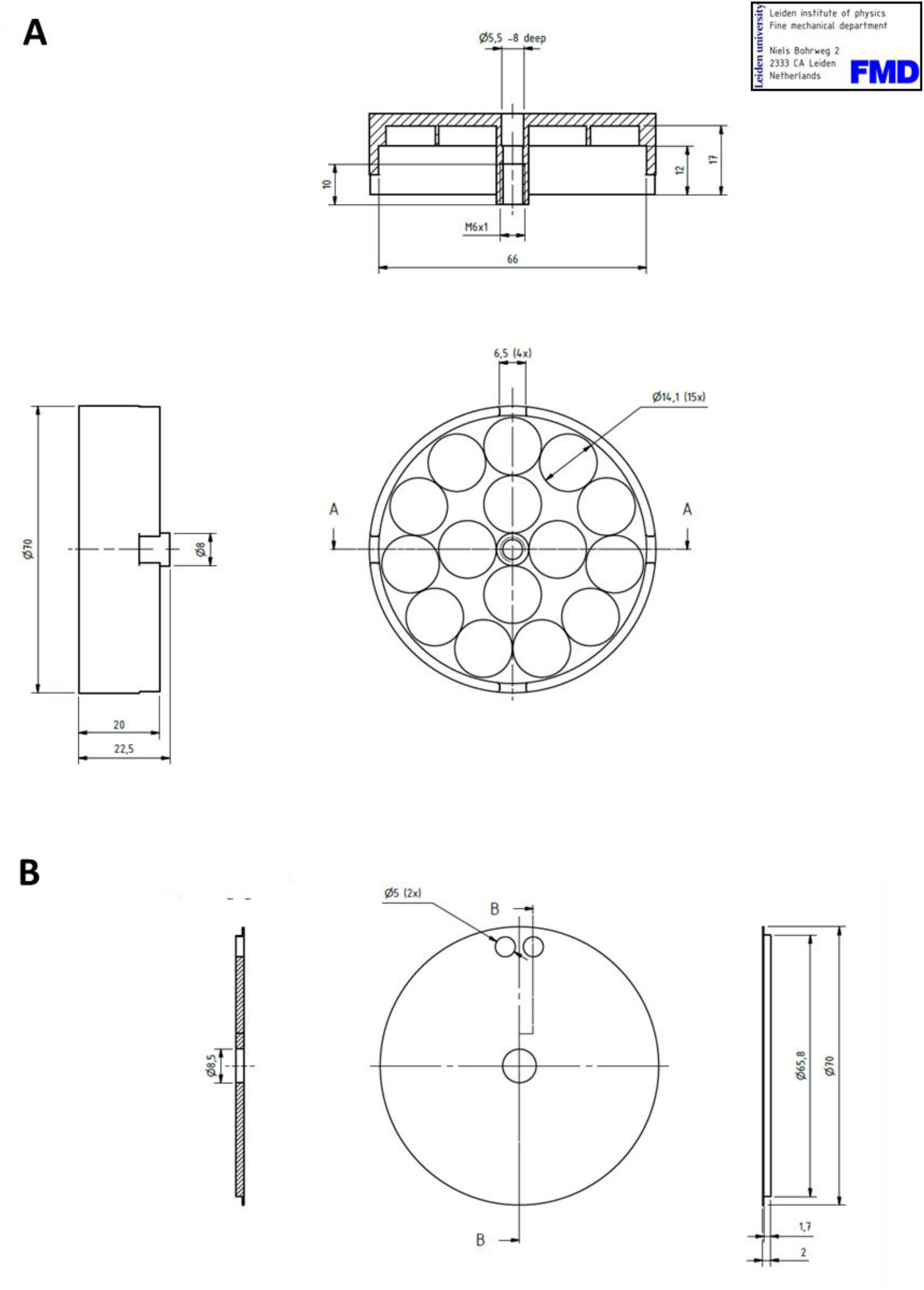
Design of the cryo-storage system. The pucks for cryogenic storage in canisters of dewar vessels. Design of both the puck (A) and the lid (B), all dimensions are depicted in millimeters.

The separate lid lies on top of the puck, held in place by both a central cylinder which is protruding 3mm over the edge of the puck (figure 1C–4) and the recessed edge of the lid (figure 1C–1). The lid is produced from 2 mm thick aluminum and has a 1.7 mm engraving in the edge, together with a central hole of 8.5 mm (figure 1C–2) to fit the central pillar. The design of the lid, combined with the higher central cylinder, makes sure the lid is well centered and offers space for any type of grid box. Two 5 mm holes in the lid facilitate its removal with a pair of tweezers (figure 1C–3). Although the lids lie firmly on the top of the pucks, they are not directly affixed to them (i.e. by magnets, threaded fittings or fasteners) to simplify the design and to minimize the risk of the lids getting stuck or frozen onto the puck. The pucks remain positioned upright as they fit tightly within the canisters.

The rod-shaped hollow cylinder in the center of the puck also serves as the connection point for manipulating the puck (figure 1C–4). This central cylinder is designed to allow easy retrieval of the puck from the canister with a 70 cm stainless-steel rod (figure D). The rod contains a spring loaded pin with a self-locking mechanism, based on a ball lock principle (M-PARTS mechanical solutions) that is controlled with a button on the rod’s handle (figure 1D). A circular plate with a diameter of 72 mm is welded onto the rod, to guide the pin to the central docking pillar of the puck while it is inside the dewar canister. The rod has four leaf-springs to facilitate the release of the puck when the button is pressed.

The storage system developed by Dr. Alasdair McDowall, uses a threaded central cylinder and a rod with matching threads at its tip to retrieve the pucks. This original rod is inserted into the dewar and turned to lock onto the puck. Our ball locked pin improves this design by minimizing the motions necessary to grasp the puck, speeds the retrieval process and decreases the chances of ice contamination interfering with the attachment.

This design requires the cryo-storage pucks to remain horizontal, in order for the retrieval-pin to align with the central cylinder. As the bottom of the canisters in our storage vessel were slightly uneven, an extra support ring was welded into the canisters to ensure that the pucks rest in the proper position.

## Work-flow using cryo-storage boxes

To transfer a cryo-storage puck, the pin of the retrieval-rod is first aligned with the central column of the puck (figure 3B). The button on the handle of the rod is then pushed to engage the self-locking pin and affix the puck to the rod (figure 3C). The puck can then be transported into the dewar for storage or to the workspace for sample manipulation (figure 3C–F). Once safely in the new location, the button is pressed again to release the puck from the rod (figure 3G–H).

**Figure 3.**
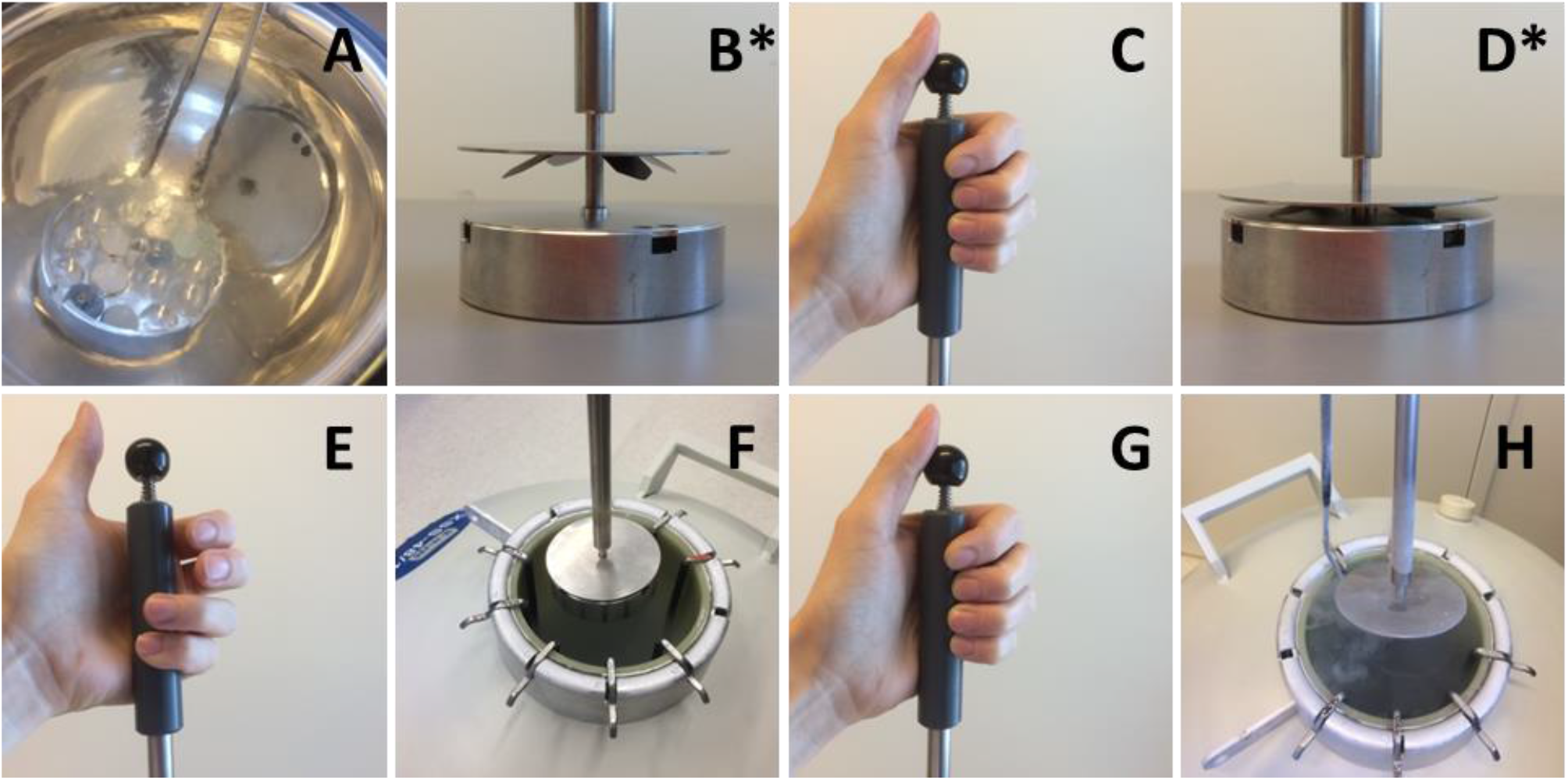
Transfer of cryo-storage pucks using the retrieval rod. The cryo-storage puck for cryo-EM grid boxes can be transferred using a retrieval rod. The cryo-EM grid boxes are positioned in the puck and covered with a lid (A). The pin of the retrieval rod is aligned with the central cylinder of the storage puck (B). The button at the end of the retrieval rod is pushed to activate the self-locking pin (C) which slides into the central cylinder and affixes the puck onto the retrieval rod (D). The pin is locked, the button can be released (E). The puck is transported to the cryogenic storage vessel (F). The puck is positioned in the target canister and released by pressing the button of the retrieval rod (G). The retrieval rod can be removed (H) and the puck is safely stored inside the dewar. Reverse the steps depicted in this figure to retrieve the puck from the storage dewar and transfer to the work space for sample manipulation. The actions of panels B^*^ and D^*^ are shown at room temperature for illustration purposes: the samples must remain submerged in liquid nitrogen continuously, as shown in panel A.

The XSS-48/10 model liquid nitrogen storage dewar (VWR systems) used to develop this design holds 10 canisters. Each canister holds 11 storage pucks and each puck can hold 15 grid boxes. As such the dewar can hold 1650 grid boxes (up to 6600 EM-grids). As all pucks will be vertically stacked upon each other, one has immediate access to the uppermost puck while the remaining pucks lie safely in liquid nitrogen storage. Pucks lower in the canister can be accessed once the top pucks have been removed: until needed the pucks are safely stored in the vessel with minimal opportunity for contamination. The canister itself can remain positioned inside the dewar completely submerged in liquid nitrogen, while transferring cryo-storage pucks.

## Other available storage solutions

The problem of inefficient and unpractical storage of cryo-EM samples has previously been addressed (Scapin, Prosise, Wismer, & Strickland, 2017). While our design was in its testing phase, several commercial solutions for cryo-EM storage were introduced. The most recent example is the SubAngstrom storage system (Scapin et al., 2017) that is based on the Rigaku ACTOR magazine system. Other designs can be purchased from MiTeGen^1^ and Molecular Dimensions^2^. Although these systems offer sophisticated and effective solutions for cryo-storage, our newly designed system is a low-cost alternative that will fit the needs of many research groups.

The design developed by our lab is easily adaptable for any type of storage dewar by adjusting the dimensions of the cryo-storage pucks. This feature offers the freedom to utilize and optimize any available storage dewar and does not require the purchase of any specific model. The inner bottom surface of the pucks was designed to hold two types of grid boxes (figure 1B). Nevertheless, the design can easily be customized to hold grid boxes with unconventional dimensions and fit the preferences of other labs. The only accessory required for this system is the steel retrieval rod. No specialized tweezer sets or canister storage shelves are required. By contracting with a local machine shop, the cost to produce the components for this system were considerably less than the price of the commercial alternatives.

As previously mentioned, the canisters themselves remain in the dewar, which we believe is an advantage over the current commercial systems which require the a whole canister to be removed from the dewar. The here presented design limits the exposure of the canisters, and consequently the other samples, to room temperature and therefore prevents excessive contamination.

We are confident that our cryo-storage system will be an effective and low cost storage solution that will be benefit the growing cryo-EM community. We will gladly provide the designs and schematics to any interested groups.

## Acknowledgements

Thanks to the Fine Mechanical Department of the Leiden University as designing and fabrication partner. The development of the cryo-storage system is funded by grant NWO BBOL 737.016.004 (Briegel) from the Netherlands Organization for Scientific Research (NWO).

We thank Mark Ladinsky for critically reading the manuscript.

1. MiTeGen http://www.mitegen.com/CEM/EMpucks/
2. Molecular Dimensions https://www.moleculardimensions.com/products/c504-Cryo-EM-Grid-box-Storage/

